# Auditory cortical alpha desynchronization prioritizes the representation of memory items during a retention period

**DOI:** 10.1101/626929

**Authors:** Nadine Kraft, Gianpaolo Demarchi, Nathan Weisz

**Affiliations:** Centre for Cognitive Neuroscience and Department of Psychology, Paris-Lodron Universität Salzburg, Austria

## Abstract

Maintaining information in working memory normally happens in dynamic environments, with a multitude of distracting events. This is particularly evident in the auditory system, for example, when trying to memorize a telephone number during ongoing background noise. How relevant to-be-memorized information is protected against the adverse influence of a temporally predictable distractor was the main goal of the present study. For this purpose we adapted a Sternberg task variant established in the visual modality, with either a strong or a weak distracting sound presented at a fixed time during the retention period. Our behavioral analysis confirmed a small, albeit significant deterioration of memory performance in the strong distractor condition. We used a time-generalized decoding approach applied to magnetoencephalography (MEG) data to investigate the extent of memory probe-related information prior to the anticipated distractor onset and found a relative decrease for the strong distractor condition. This effect was paralleled by a pre-distractor alpha power decrease in the left superior temporal gyrus (STG), a cortical region putatively holding memory content relevant information. Based on gating frameworks of alpha oscillations, these results could be interpreted as a failed inhibition of an anticipated strong (more salient) distractor. However, in a critical analysis we found that reduced alpha power in the left STG was associated with relatively increased memory probe-related information. Our results therefore support the view of alpha power reductions in relevant sensory (here auditory) cortical areas to be a mechanism by which to-be-remembered information is prioritized during working memory retention periods.

## Introduction

Adaptive sensory processing entails the prioritization of task-relevant features with respect to competing information. Top-down modulation of activity in neural ensembles encoding task-relevant or distracting information respectively are crucial in achieving this goal. Alpha oscillations have been linked to such a putative top-down mediated gain modulation, with enhanced alpha power marking relatively inhibited states (Jensen and Mazaheri, 2010; Klimesch et al., 2007). Especially for the visual modality, a vast amount of empirical evidence supports this notion, for example, increased alpha power in parieto-occipital cortical regions contralateral to the unattended hemifield is a very robust finding (e.g. (Busch and VanRullen, 2010; Thut, 2006)); when attending to the auditory modality while ignoring upcoming distracting visual input alpha power is enhanced in parieto-occipital cortical regions (Frey et al., 2014; Fu et al., 2001; Snyder and Foxe, 2010); the general inhibitory gating function of localized alpha increases has also been reported with respect to more specific visual features (Jokisch and Jensen, 2007; Zumer et al., 2014). Also for the domain of working memory, alpha increases have been reported during the retention period in the visual (e.g. (Jensen et al., 2002; Klimesch et al., 1999)), somatosensory (e.g. (Haegens et al., 2009)) and auditory modality (e.g (Obleser et al., 2012)), putatively protecting the to-be-remembered information against interference. While this load dependent top-down amplification of alpha is widely accepted, also circumscribed decreases in alpha power (often labeled as desynchronization) have been deemed functionally important in the context of working memory tasks, reflecting an enhanced activation of performance-relevant neural ensembles (e.g. (Noh et al., 2014; Sauseng et al., 2009); see (van Ede, 2018) for review). A recent framework by Hanslmayr et al. aiming at explaining the role of neural oscillations underlying episodic memory (Hanslmayr et al., 2016) explicitly links the extent of alpha desynchronization to the representational strength of the information content. This is in line with a framework by van Ede (van Ede, 2018) who stresses the importance of circumscribed alpha decreases when item-specific information needs to be prioritized in the retention period of working memory tasks. In order to exert optimal control over perceptual processing, the brain exploits relevant cues such as temporal regularities (Rohenkohl and Nobre, 2011; van Ede et al., 2018) to regulate the excitatory-inhibitory balance in (ir-)relevant neural ensembles in an anticipatory manner.

In this study we are interested in scrutinizing the aforementioned issue in the auditory system. Distracting sounds are ever-present in natural listening environments; therefore flexible inhibition of distracting and strengthened representations of relevant sounds has to be provided during auditory processing. Interestingly, an increasing amount of evidence points to a functional role of alpha oscillations in listening tasks - for example, selective attention or memory - similar to other sensory modalities (Frey et al., 2015; Weisz and Obleser, 2014). As mentioned above, increases of alpha have been observed over posterior brain regions when focussing attention on auditory input, a pattern also observed in challenging listening situations, for example, with increased cognitive load or when faced with background noise (for reviews see (Johnsrude and Rodd, 2016; Rönnberg et al., 2011)). However, increases of alpha as a mechanism for selective inhibition (Strauß et al., 2014) have rarely been shown for auditory cortex. With regards to alpha desynchronization, different lines of evidence showing an association between (also illusory) sound perception with low auditory cortical alpha power (e.g. (Lange et al., 2014; Weisz et al., 2007; Weisz and Obleser, 2014)), suggest a link to representational content as described above (Hanslmayr et al., 2016). The goal of the present study was to test whether auditory cortical alpha oscillations are modulated in an anticipatory manner prior to an upcoming distractor presented in the same modality. In particular we were interested in whether such putative alpha modulations would follow more an “inhibition” (auditory cortical alpha increases expected) or “prioritization” (auditory cortical alpha decreases expected) account. Furthermore we wanted to test the relationship between such anticipatory alpha modulations and the extent to which task-relevant auditory information is protected from interference.

To investigate the outlined issues we adapted a Sternberg task variant introduced by Bonnefond and Jensen (Bonnefond and Jensen, 2012) to the auditory modality. These researchers illustrated pronounced alpha increases as well as phase effects in parieto-occipital regions, prior to the presentation of a more potent but temporally predictable visual distractor presented in the retention period. Using magnetoencephalography (MEG), our study shows marked differences for an analogous task in the auditory system, with alpha desynchronization prior to the strong distractor in left auditory cortical regions putatively relevant for the representation of the verbal memory probe. Importantly, using multivariate pattern analysis (MVPA) we show that lower pre-distractor alpha power in left auditory cortex goes along with enhanced representation of the memory probe during the retention interval. Overall our study supports the important role of circumscribed alpha decreases in prioritizing relevant information in working memory (van Ede, 2018).

## Results

Thirty-three healthy participants performed a modified Sternberg task (Bonnefond and Jensen, 2012) adapted to the auditory modality. In each trial they listened to a sequence of four consonants spoken by a female voice (see Figure 1A). These items had to be memorized across a 2 s retention period, after which a target item was presented. Participants were requested to report whether the target item was part of the memory set or not. Critically, following precisely 2 s after onset of the last memory item, a distractor was presented which depending on the block was either a consonant spoken by a male voice (i.e. strong distractor) or a scrambled letter (i.e. weak distractor).

**Figure 1:**
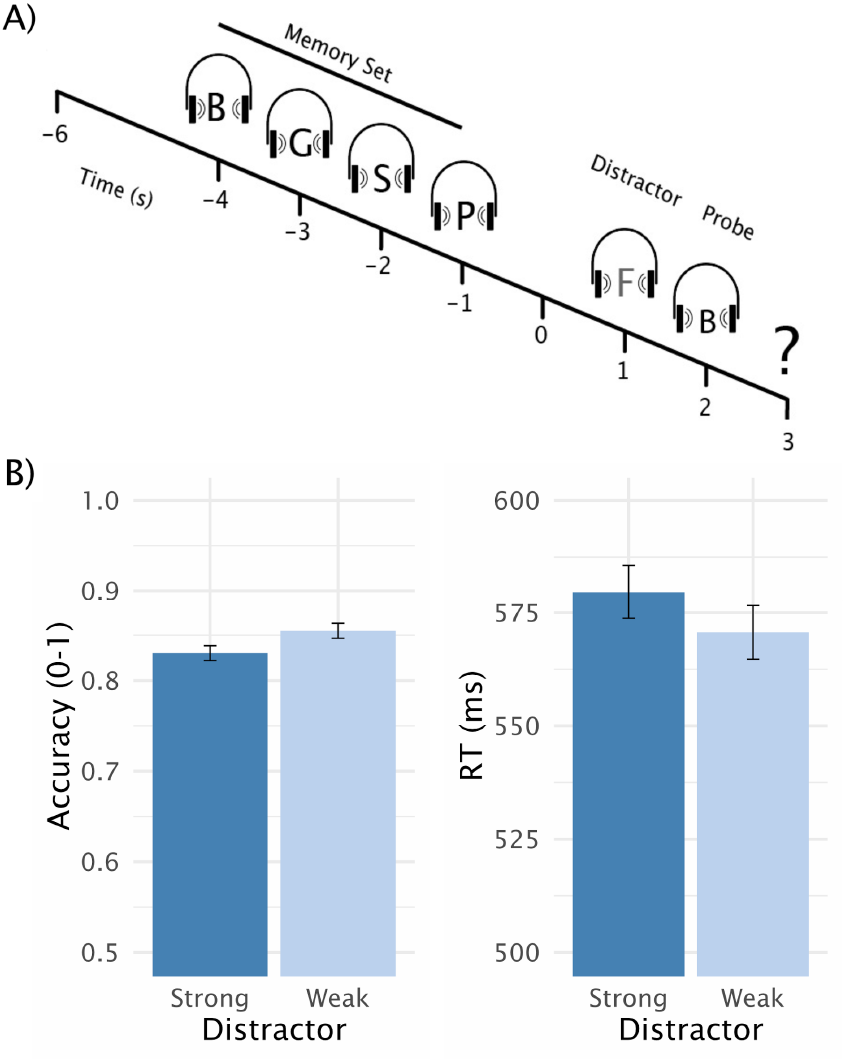
The Modified Auditory Sternberg Paradigm A) A sequence of four consonants spoken by a female voice was presented. After the retention period either a strong (consonant spoken by a male voice) or a weak (scrambled consonant) distractor was presented (at 1 s). Subsequently participants indicated by a button press whether the probe was part of the memory set (“part”) or not (“no part”). B) Accuracy and reaction times (RT) responding to the required task of deciding whether the probe was part of the memory set. Although the RT were independent from the distractor type (right panel), the accuracy was significantly lower for strong versus weak distractors (*t*_32_ = −2.11, *p*_one-sided_ = .02), indicating a more challenging strong distractor condition (right panel).

### Adverse behavioral impact of strong distractors

We reasoned that the processing of a strong distractor would be more difficult to suppress and should affect behavioral performance. Comparing average accuracy between the strong and the weak distractor conditions showed a small (85% vs. 83%) but statistically significant (*t*_32_ = −2.11, *p*_one-sided_ = .02) deterioration of performance for strong distractors (Figure 1B). Reaction times were on average 9 ms slower for the strong distractor condition (579 ms vs. 570 ms); however, this difference was not significant (*t*_29_ = 1.07, *p*_one-sided_ = .14). It should be noted that speed was not emphasised for responses to avoid interference from button presses on processing of the target item. This may have reduced potential reaction time differences. Overall, the behavioral analysis supports the notion that the strong distractor condition was slightly more challenging, laying a solid foundation for the subsequent MEG analysis.

### Decoding probe-related information

Our main goal was to investigate to what extent different distractor levels influence the strength of memory probe-related information during the retention period, in particular prior to the predictable onset of the distractor. Also, we wanted to relate these effects to potential alpha power modulations in the pre-distractor period. To this end we first applied temporal decoding, using linear discriminant analysis (LDA; see *Materials and Methods*), on the post-target MEG sensor-level activity, to classify whether a target was part of the memory set or not. The results are depicted in Figure 2A showing robust and sustained above-chance classification performance commencing ~334 ms after target onset (*p*_cluster_ = 4e-4) and lasting until the onset of the response prompt at 700 ms post-target. In subsequent analyses this post-target period decoding was used as training set and the derived classifiers were applied in a time-generalized manner to the preceding retention period (see below). While the results so far show that probe relevant information can be differentiated based on the MEG data, the resulting temporal decoding pattern uses all sensors and is therefore spatially agnostic. In order to obtain insights into which brain regions may be contributing to the effect, we adapted an approach to derive informative activity in source space (Marti and Dehaene, 2017). In brief, this approach projects the sensor level classifier weights to source space using beamformer filters. To make the effects more interpretable, we implemented a within-subject permutation analysis and z-scored the classifier-weights in a first-level analysis. These were subsequently tested on a group-level against zero within a nonparametric cluster permutation test using a *t*-test. As shown as an inset of Figure 2A, informative activity related to probe relevant information can be detected in widespread cortical regions encompassing temporal, parietal and frontal areas. While a bilateral pattern can be observed, there was a clear left-hemispheric dominance with the most pronounced effect localizable to left superior temporal gyrus (STG). Given the particular involvement of this latter area in processing speech sounds (e.g. (Mesgarani et al., 2014)), it was used as a region of interest for the spectral analysis (see below).

**Figure 2:**
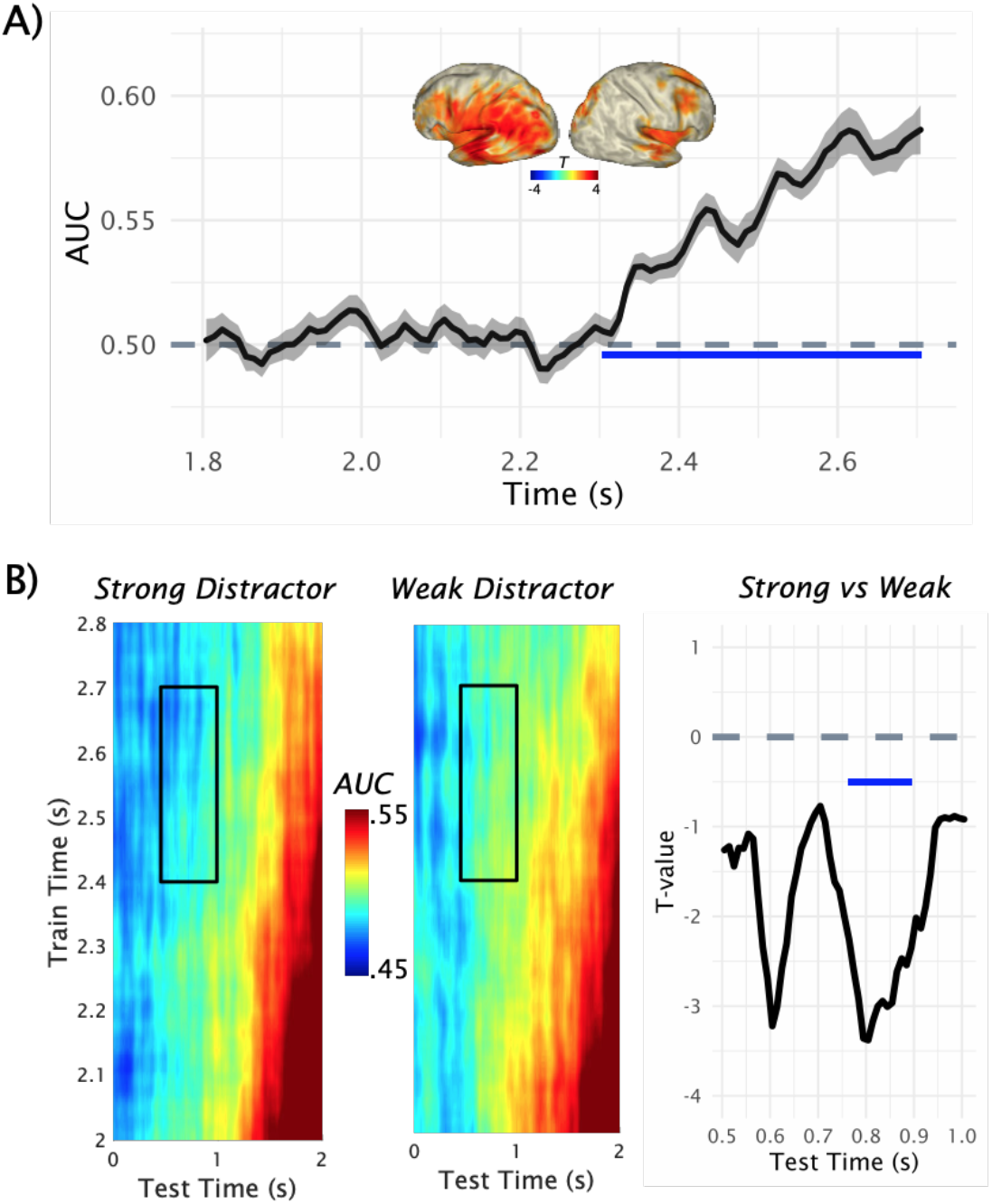
The Decoding of Probe-Related Information. A) Results of the temporal decoding on the MEG sensor-level activity after the target presentation to classify whether a target was part of the memory set or not. Above chance detection performance (AUC = area under the ROC curve) was found commencing ~300 ms after the target onset (at 2.0 s) and lasting at least until the response was prompted. Activity related to target relevant information can be seen in cortical regions with a clear left-hemispheric dominance. B) The time generalization result is shown separately for the strong and weak distractor condition (left and middle panel). The strongest classification results are obtained approximately at the onset of the target (at 2 s). Relatively decreased decoding performance was obtained prior to the onset of the strong distractor. Statistical comparison of strong vs. weak distractor conditions revealed two peak effects at ~400 ms and ~200 ms preceding the distractor onset, although only the effect closer to distractor onset was significant (*p*_cluster_ = .0156) at a cluster level (right panel).

After illustrating the above-chance decoding performance for probe-related information, we tested to what extent this pattern is present in the retention period and how it is modified by the distractor especially prior to its anticipated onset. The classifier described above was employed for this purpose using time generalization (King and Dehaene, 2014). The full time generalization result is shown separately for the strong and weak distractor condition in Figure 2B. As an aside, it can be seen that the strongest classification results are obtained approximately at the onset of the target (at 2 s), which corresponds to the area close to the diagonal of the time generalization matrix. We were mainly interested in the decoding performance in the period preceding the anticipated distractor at 1 s, that is, in the off-diagonal pattern. On a descriptive level, decreased decoding performance was obtained prior to the onset of the strong distractor. Statistical analysis was done for a 500 ms window preceding the onset of the distractor (Figure 2B, right panel), focussing on a 400-700 ms training time window (see above). This analysis yielded two peak effects at ~400 ms and ~200 ms preceding the distractor onset during which critical *t*-values (±2.0369) were exceeded. However, only the latter cluster was significant following a nonparametric permutation test (*p*_cluster_ = .0156). These results suggest that probe-related information is differentially activated prior to distractor onset, with relatively reduced activation prior to the strong distractor. This effect is in line with the behavioral results described above, implying an overall detrimental impact of a strong distractor.

### Pre-distractor alpha modulations

As the next step, we focussed on alpha power modulations in the left STG, with an emphasis on the period immediately preceding the predictable occurence of the distractor. In the case that this task relevant auditory processing region did prepare to inhibit processing of this irrelevant sound, alpha enhancements would be expected in particular preceding the strong distractor. The time-frequency representations in Figure 3A, displaying the induced power in the 5-25 Hz range, show strong ongoing alpha / beta activity with a peak ~10 Hz in the left STG. A 500 ms period preceding the occurrence of the distractor is marked, suggesting an alpha power decrease in the strong as compared with the weak distractor condition. This impression is supported by a nonparametric permutation test (*p*_cluster_ = .0104), yielding a significant difference in this period over an alpha to beta range with a maximum difference ~12 Hz (Figure 3A, right panel). Given the strong temporal predictability of the distractor occurrence, stronger prestimulus phase alignment of alpha oscillations could be expected as was reported in the visual modality by Bonnefond and Jensen (Bonnefond and Jensen, 2012) (see however (van Diepen et al., 2015)). This process putatively exploits the fact that excitability varies over an alpha cycle, to optimally align its inhibitory phase to maximally suppress processing of the irrelevant sound. However, even though clear post-distractor evoked alpha enhancements could be observed (see Figure 3B), no prominent evoked alpha could be observed preceding the distractor. For the sake of completeness, we ran an analogous statistical test as for the induced power, showing no difference at the cluster corrected level (Figure 3B, right panel). Since no pronounced evoked alpha activity was identified in the pre-distractor period, we refrained from further analysis (such as phase opposition effects, etc.). This result extends a previous report (van Diepen et al., 2015) in finding no evidence that auditory cortical alpha phase is adjusted in a top-down manner.

**Figure 3:**
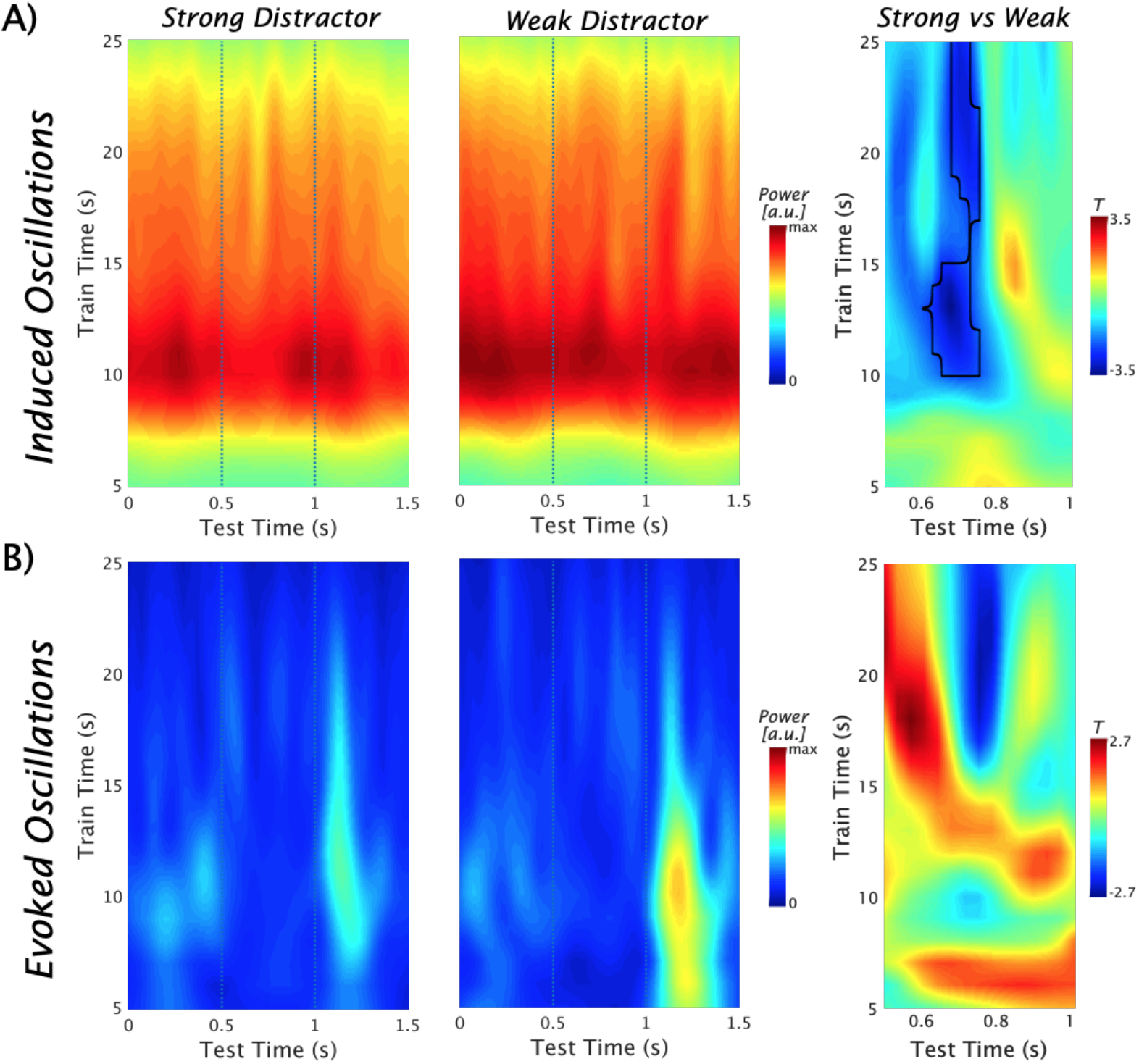
The Pre-distractor Alpha Power Modulations in the Left Superior Temporal Gyrus. A) Time-frequency representations of the induced power show strong ongoing alpha / beta activity with a peak ~10 Hz. The vertical dots indicate a 500 ms period preceding the occurrence of the distractor. An alpha power decrease in the strong vs. the the weak distractor condition can be seen (left and middle panel). The notion is supported by the outcome of a nonparametric permutation test leading to a significant difference at cluster level (marked by a black contour; *p*_cluster_ = .0104) over an alpha to beta range with a maximum difference ~12 Hz (right panel). B) Time-frequency representations of the evoked power. Post-distractor alpha enhancements are seen, but no prominent alpha preceding the distractor (left and middle panel). The nonparametric statistical test at cluster level showed no difference (right panel).

Overall, the spectral analysis described in this part suggests a pronounced alpha decrease prior to the expected occurrence of the strong distractor. Taken together with the behavioral data as well as the decoding result, this alpha decrease could be interpreted as dysfunctional, which would also fit with views emphasizing the beneficial role of alpha enhancements in listening tasks (Strauß et al., 2014). For example, the expectation of a more salient auditory distractor may involuntarily draw more selective attention towards it, making it more difficult to suppress. As an alternative to this failed inhibitory gating view, it is also possible that memory probe-related information are prioritized to protect the representation against the detrimental effect of the anticipated strong distractor (van Ede, 2018). Based on results presented so far, these alternatives cannot be differentiated. In the next part, we will attempt to address this important issue by linking pre-distractor alpha power modulations to the presence of probe-related information.

### Pre-distractor alpha power modulations of probe-related information

To address the functional relevance of pre-distractor alpha power modulations in the left STG in greater detail, trials were sorted according to alpha power (10-16 Hz) in this region in a 400-1000 ms time period following the onset of the retention period (i.e. a 600 ms pre-distractor window). Subsequently, these trials were median split into a high and low alpha power bin. Analogous to the analysis described above (see also Figure 2), we trained a classifier on all trials to discriminate whether a probe was part of a memory set or not and applied the classifier to a +/− 500 ms time-window around the distractor presentation separately to the high and low alpha trials. Based on the previous analysis we focussed on a 400-700 ms training time period and calculated the difference in decoding accuracy between high and low alpha trials for each individual participant. In a next step we (Pearson) correlated this difference time series with the individual *log10* alpha power ratio between high and low alpha. Nonparametric cluster permutation test revealed a significant negative correlation ~170-90 ms (*p*_cluster_ = .0272) prior to distractor onset and another period ~180-500 ms (*p*_cluster_ = .0012) following the onset of the distractor (see Figure 4A). To gain an impression of these effects, scatterplots for relevant pre- and post-distractor time-points are displayed in Figure 4B, showing that the effects are not driven, for example, by outliers. A negative correlation in this case means that across subjects, decoding of probe-related information in the retention period deteriorated with stronger alpha power in the left STG. While carefully balancing the frequency of strong and weak distractor trials in both alpha power bins, we wanted to ensure that the described effect is indeed related to the decoding of probe-related information and not driven by the condition differences described above. We thus repeated the analysis training the classifier to decode the experimental condition and applied the same training time windows to the testing period around the distractor presentation again for high and low alpha trials. The same correlation approach shows an absence of effects for this analysis, ensuring that the aforementioned effects are specific to probe-related information in the retention period. Overall, the analysis presented here suggests that alpha power reductions prior to the presentation of strong distractors could be functionally important in activating memory probe-related information rather than being a dysfunctional automatic orientation to the strong distractor.

**Figure 4:**
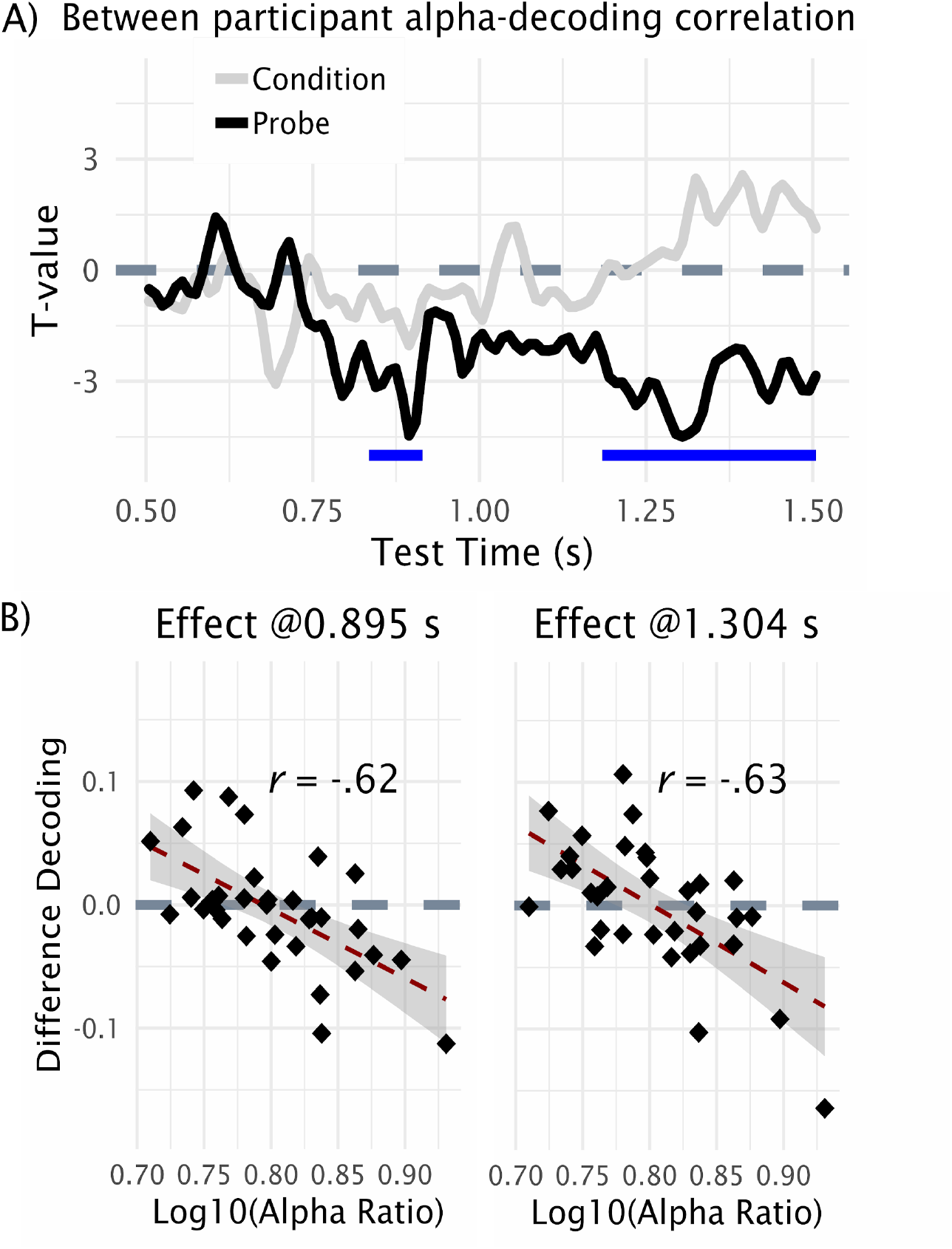
Correlation between alpha power modulation and memory probe-related information in pre- and post-distractor period. A) Magnitude of alpha power changes and memory probe-related information was computed within-participant and correlated across the group. Results reveal a negative relationship (black line), which was not observed when running the same analysis using condition-related information for the correlation (grey line). B) Scatterplots for the two time-periods marked in A), showing that effects are not driven by outliers.

## Discussion

In the current study, we investigated the neural dynamics prior to an anticipated distractor in the auditory modality. We were particularly interested in potential modulations of alpha power in auditory cortical regions, which have shown similar patterns as described in various cognitive tasks in the visual system (Frey et al., 2015). Also alpha power modulations, although mainly in non-auditory regions, have been previously linked to listening effort or attentional control (McGarrigle et al., 2014; Pichora-Fuller et al., 2016; Wöstmann et al., 2017). In order to understand the functional relevance of potential alpha power modulations, it was important to link them to the informational content carried by the neural patterns in the same time period. To achieve our goals we adapted a modified Sternberg paradigm first proposed by Bonnefond and Jensen (Bonnefond and Jensen, 2012) to the auditory system: that is, we introduced - in a blockwise manner - putatively weak and strong auditory distractor items in the retention period with a predictable timing. While behavioral effects appear overall somewhat weaker than in the original visual experiment by Bonnefond and Jensen, lower accuracy was obtained for strong distractors. The behavioral finding suggests an adverse effect of strong acoustic distractors on the representation of probe-related information. This notion was supported on a neural level using a time-generalization approach (King and Dehaene, 2014), in which classifiers were trained to decode whether a probe item was part of the memory set or not and subsequently applied to the period around the distractor presentation in the retention interval. We could indeed show weaker probe-related information already in the anticipatory period of the distractor. Interestingly alpha power (extending into the beta frequency range) prior to the presentation of the strong distractor decreased, whereas it was relatively sustained in the weak distractor condition. This effect is at odds with findings reported in the visual modality using an analogous paradigm (Bonnefond and Jensen, 2012), where - in line with ideas of an inhibitory role of alpha oscillations (Jensen and Mazaheri, 2010; Klimesch et al., 2007) - induced alpha enhancements were seen prior to a strong distractor. Also in the Bonnefond and Jensen study pronounced pre-distractor evoked alpha power effects were observed that could not be identified in our auditory task. An integration of our alpha findings with these lines of evidence appear challenging based on power effects alone, as they could be interpreted either as involuntary direction of attentional resources to the anticipated strong distractor (failed inhibition within a gating account) or as a top-down driven prioritization of probe-related information in anticipation of a strong distractor (see e.g. (Hanslmayr et al., 2016)). The group-level effect of reduced decoding accuracy prior to the strong distractor could be seen as a support for the first interpretation, yet this result does not take into account the interindividual variability of the decoding and alpha power effects. Indeed across participants we could observe a strong correlation between the modulation of alpha power and changes in decoding accuracy, with strong relative pre-distractor alpha enhancements going along with reduced decoding accuracy (also in the post-distractor period). This result suggests decreases of alpha power in the left auditory cortex are linked to a prioritization of memory probe-related information, such that anticipatory alpha reductions prior to the strong distractor are adaptive in terms of the task.

While differences to the original visual experiment (Bonnefond and Jensen, 2012) may point to modality specific processes, our results may also seem at odds with findings for the auditory modality which frequently point to alpha enhancements as an adaptive process within challenging listening tasks (for review see (Strauß et al., 2014)). In line with dominant views regarding functional relevance of alpha oscillations (Jensen and Mazaheri, 2010; Klimesch et al., 2007), alpha enhancements in such circumstances have been linked to selective inhibition of irrelevant “channels” of auditory information (Strauß et al., 2014). However, the sources of these alpha enhancement effects have most frequently been identified in non-auditory brain regions (e.g. (Obleser and Weisz, 2012; Wilsch et al., 2015)) and rarely in the auditory cortex (e.g. (Müller and Weisz, 2012)). Indeed reduction of auditory cortical alpha activity has been commonly linked to attended (Frey et al., 2014) as well as perceived (including illusory) auditory input (e.g. (Leske et al., 2014; Müller et al., 2013; Weisz et al., 2007); for a more general perspective see (Lange et al., 2014)). However, the association of alpha modulations to attended / ignored or perceived auditory information has so far been very indirect. Our study significantly advances this state by showing an inverse relationship between alpha power modulations in the left auditory cortex and memory probe-related information in the retention period. This result supports the aforementioned interpretations of studies showing alpha power reductions in the auditory modality and a more general assertion of cortical alpha desynchronization during memory tasks to represent the content of memorized information (Hanslmayr et al., 2016). Overall, we find that our results can be reconciled with the previous studies focussing on alpha enhancements, as alpha reductions or enhancements may show engagements of different neural systems in processing relevant or blocking irrelevant information respectively (van Ede, 2018). The functional versatility of alpha power modulations in listening tasks also serves as a precaution not to simplistically equate, for example, alpha power enhancements to concepts such as listening effort (McGarrigle et al., 2014; Pichora-Fuller et al., 2016).

In summary, precise predictability of the occurrence of an auditory distractor leads to an anticipatory prioritization of memory probe-related information. We show that modulations of alpha oscillations in task-relevant auditory cortical regions could be a relevant process mediating the “protection” of relevant auditory information against interference. In doing so, our study significantly adds to our understanding of the functional role of alpha oscillations in the auditory system.

## Materials and Methods

### Participants

Thirty-three participants were included in the calculations (22 female; age range: 18-46 years; mean age: 26.8 years). Four participants were excluded due to technical issues during the testing or because the data were too noisy. All participants reported normal or corrected-to-normal vision and an absence of hearing problems in daily life. None of them suffered or was suffering a psychological or neurological disorder. Written informed consent was obtained from each participant prior to the experiment. They obtained either €10/h reimbursement or credits required for their bachelor studies in psychology. All procedures were approved by the Ethics Committee of the University of Salzburg.

### Stimuli and Procedure

Participants underwent standard preparation procedures for MEG experiments. Five head position indicators (HPI) coils were applied (three on the forehead, and one behind each ear). Using a Polhemus FASTRAK digitizer, anatomical landmarks (nasion, left and right pre-auricular points) and HPI coils were recorded, and additionally approximately 300 head shape points were sampled. To control for eye movements and heart rate, electrodes were applied horizontally and vertically to the eyes (electrooculogram), one electrode was placed on the lower left ribs and one next to the right clavicle (electrocardiogram), as well as one reference electrode on the back. After entering the MEG cabin, a five min resting state was recorded, which was not utilized for the present study. The experimental paradigm consisted of a Sternberg task, similar to the one used by Bonnefond and Jensen (Bonnefond and Jensen, 2012), but adapted to the auditory modality. Visual stimuli were displayed with the PROPixx projector (VPixx Technologies Inc.) on a opaque screen. Auditory stimuli were delivered using the SOUNDPixx system (VPixx Technologies Inc.) through two pneumatic tubes. The stimulus delay introduced by the tube was measured using a microphone (16.5 ms ± 0.1 ms), and this delay was taken into account and compensated in the analysis phase. The experiment was programmed in MATLAB 9.1 (The MathWorks, Natick, Massachusetts, U.S.A) using the open source Psychophysics Toolbox [40].

During the experiment participants focused on a fixation point. They listened to a memory set of four consonants spoken by a female voice (see Figure 1A). The interstimulus interval between the consonants presentation onsets was set to 1 s, and a distractor was presented to the participants 2 s after the final fourth letter (timepoint 1 s in Figure 1A). Within each experimental block, the distractor was either a consonant spoken by a male voice (strong distractor) or a temporally scrambled consonant (weak distractor). The scrambling of the distractor was achieved using the Matlab-based *shufflewins* function (Ellis, 2011). This scrambling approach preserves the frequency content of the original voice but makes it unintelligible. One second after the distractor, the probe was presented spoken by the same female voice as in the memory set. Thereafter the participants needed to decide via button press whether the probe was part of the memory set or not. The participants were exposed to 12 blocks, 6 per each distractor condition and each one containing 24 trials. An intertrial interval from 1.5-2.5 s (mean 2.0 s, uniformly distributed) was used. One block had a duration of about 6 min. The sequence of the conditions and the assignment of the buttons was randomized across participants.

### MEG acquisition and analysis

The brain magnetic signal was recorded (sampling rate: 1 kHz, hardware filters: 0.1 - 330 Hz) using a whole head MEG device (Elekta Neuromag Triux, Elekta Oy, Finland) in a standard passive magnetically shielded room (AK3b, Vacuumschmelze, Germany). Signals were captured by 102 magnetometers and 204 orthogonally placed planar gradiometers at 102 different positions. We used a signal space separation algorithm (Taulu and Kajola, 2005) implemented in the Maxfilter program (version 2.2.15) provided by the MEG manufacturer to remove external noise from the MEG signal (mainly 16.6 Hz, i.e. Austrian train AC power supply frequency, and 50 Hz plus harmonics) and realign data to a common standard head position (to [0 0 −4] cm, *-trans default* Maxfilter parameter) across different blocks based on the measured head position at the beginning of each block. First, a high-pass filter at 0.5 Hz (6th order zero-phase Butterworth filter) was applied to the continuous data. Then, continuous data were epoched around the onset of the retention phase using a 3 s pre- and post-stimulus window. For most analyses, the data were downsampled to 256 Hz (100 Hz for decoding analysis; see below). The epoched data were subjected to an Independent Component Analysis using the runica algorithm (Delorme and Makeig, 2004). The components were manually scrutinized to identify eye blinks, eye movements and heartbeat, resulting in approximately two to five components that were removed from the data. Given this extensive preprocessing, no trials had to be rejected.

In a first step we applied an LDA classifier to a time window -.2 to .7 s centered on the probe presentation to confirm that by using all MEG sensors and all trials we could decode whether a probe was part of the four-item memory set or not. Apart from using the classifier weights to identify areas containing informative activity (see below), the trained classifier was applied in a time-generalized manner (King and Dehaene, 2014) to the retention period separately for the strong and weak distractor condition. By focussing in particular on the .5 s period prior to the presentation of the distractor and a training time period (.4-.7 s) in which the classifier showed above-chance performance, we could test to what extent memory probe-related information was modulated in anticipation of the distracting sound. Given the nature of our research question outlined in the introduction, we wanted to analyze pre-distractor alpha power modulations in memory probe-relevant brain regions. For this purpose, covariance-corrected (Haufe et al., 2014) classifier weights were projected to source space using an approach adapted from Marti and Dehaene (Marti and Dehaene, 2017). A realistically-shaped single-shell head model (Nolte, 2003) was computed by warping a template MNI brain to the participant’s head shape. A grid with 1 cm resolution on the template brain was morphed to fit the individual brain volume and lead fields were computed for each grid point. This information was used along with the covariance matrix of all sensors computed via the entire 30 Hz low-pass filtered epoch to obtain LCMV spatial filters (Van Veen et al., 1997). These beamformer filters were subsequently multiplied with the aforementioned covariance-corrected classifier weights to obtain “informative activity” (Marti and Dehaene, 2017) in source space (taking the absolute value on source level). In order to make this data more interpretable we implemented a permutation approach converting these time series to z-values and testing them across participants against 0 (see below). Overall this data-driven approach yielded meaningful neuroanatomical regions differentiating whether a probe was part of the memory set or not. Given the particular interest in auditory processes we focussed on the left STG which was the region providing the most prominent informative activity. For this region we used the beamformer filters to project the single trial data onto a left STG virtual sensor and applied spectral analysis on it. More precisely we used Fourier transform of Hanning-tapered data applied to a frequency range of 2-30 Hz (in 1 Hz steps) and time shifted between a period of −1.5 to .5 s around onset of the distractor (shifted in steps of .025 s). The time window for the spectral analysis was adapted to each frequency (4 cycles) and the analysis was performed separately for the strong and weak distractor condition.

Data preprocessing, spectral and source analysis was done using the Fieldtrip toolbox (Oostenveld et al., 2011). For the decoding analysis we used the Matlab based open-source MVPA-Light toolbox (https://github.com/treder/MVPA-Light).

### Statistical analysis

The behavioral impact of the distractor types was tested using a paired t-test, comparing accuracies and reaction times. Given the hypothesis that the strong distractor would be detrimental to performance, one-tailed testing was performed. With regard to our trained classified decoding, accuracy was tested against chance level (AUC = .5) between -.2 to .7 s around probe onset using a t-test. In order to make the source projected classifier weights (“informative activity”) more interpretable, we generated randomly shuffled trial labels and reran the same classifier and source projection approach. This was done 500 times and the empirically observed values at each time and grid point were z-transformed using the mean and standard-deviation from the randomized data. The z-transformed data were tested against 0 across participants using a t-test. Also the time-generalized decoding analysis and the spectral analysis described above were assessed using a t-test comparing the strong vs. weak distractor condition. To control for multiple comparisons we employed a nonparametric cluster permutation approach as proposed by Maris and Oostenveld (Maris and Oostenveld, 2007) normally using 5000 randomizations. Finally, to test the relationship of within-subject alpha modulations and memory probe-related information in the distractor period, we first binned the trials in high and low alpha (13 Hz according to maximum condition effect) trials according to a .6 s pre-distractor time window of our left STG region of interest. We applied the trained classifier to the retention periods of these binned trials separately and subsequently subtracted these time-generalized decoding accuracies (averaged over a training time period of 2.4-2.7 s; see Figure 2A). The relationship of alpha modulation (binned high vs. low alpha trials) and modulations of memory probe-related information was tested using a Pearson correlation. A nonparametric permutation test was used as described previously focussing on a period of -.5 to .5 s with respect to the distractor onset. In order to strengthen the interpretation that relevant results can be attributed to probe-relevant information, an analogous approach was performed applying a classifier trained to decode the conditions (i.e. strong vs. weak distractor) from the analogous 2.4-2.7 s time period following probe onset.

## Acknowledgements

We would like to thank Jens Gfroerer-Kötschau and Manfred Seifter for their support during data collection.

